# Posterodorsal medial amygdala urocortin-3, GABA and glutamate mediate suppression of LH pulsatility in female mice

**DOI:** 10.1101/2022.07.07.499104

**Authors:** Deyana Ivanova, Xiao-Feng Li, Caitlin McIntyre, Kevin T O’Byrne

## Abstract

The posterodorsal subnucleus of the medial amygdala (MePD) is an upstream modulator of the hypothalamic-pituitary-gonadal (HPG) and hypothalamic-pituitary-adrenal (HPA) axes. Inhibition of MePD urocortin-3 (Ucn3) neurons prevents psychological stress-induced suppression of LH pulsatility while blocking the stress-induced elevations in corticosterone (CORT) secretion in female mice. We explore the neurotransmission and neural circuitry suppressing the GnRH pulse generator by MePD Ucn3 neurons and we further investigate whether MePD Ucn3 efferent projections to the PVN control CORT secretion and LH pulsatility. Ucn3-cre-tdTomato female ovariectomised (OVX) mice were unilaterally injected with AAV-ChR2 and implanted with optofluid cannulae targeting the MePD. We optically activated Ucn3 neurons in the MePD with blue light at 10Hz and monitored the effect on LH pulses. Next, we combined optogenetic stimulation of MePD Ucn3 neurons with pharmacological antagonism of GABAA or GABAB receptors with bicuculline or CGP, respectively, as well as a combination of NMDA and AMPA receptor antagonists, AP5 and CNQX respectively, and observed the effect on pulsatile LH secretion. A separate group of Ucn3-cre-tdTomato OVX mice with 17ß-estradiol (E_2_) replacement were unilaterally injected with AAV-ChR2 in the MePD and implanted with fibreoptic cannulae targeting the PVN. We optically stimulated the MePD Ucn3 efferent projections in the PVN with blue light at 20Hz and monitored the effect on CORT secretion and LH pulses. We reveal for the first time that activation of Ucn3 neurons in the MePD inhibits GnRH pulse generator frequency via GABA and glutamate signalling within the MePD, while MePD Ucn3 projections to the PVN modulate the HPG and HPA axes.

## Introduction

Stress exerts a profound suppressive effect on reproduction in mammals, including rodents and humans. The amygdala, part of the limbic system, is a key emotional centre integrating incoming cues from the external environment, such as anxiogenic signals, with the reproductive and stress axes (1). The posterodorsal sub-nucleus of the medial amygdala (MePD) is an upstream modulator of pulsatile luteinizing hormone (LH) secretion (2–4) and exerts an inhibitory break on pubertal timing (5,6). The MePD sends direct projections, of an unknown phenotype, to the kisspeptin (kiss1) neuronal population in the hypothalamic arcuate nucleus (ARC) (7,8) known as KNDy because they co-express neurokinin B (NKB) and dynorphin A (Dyn) (9–11). The synchronous activity of the ARC KNDy network drives pulsatile gonadotropin-releasing hormone (GnRH) and LH release (11–14). The MePD also contains a kiss1 neuronal population and selective optogenetic activation of these neurons increases GnRH pulse generator frequency in mice (4) and unpublished observations show this is an effect involving both MePD GABA and glutamate signalling.

We have recently shown that urocortin-3 (Ucn3), a member of the corticotropin-releasing factor (CRF) stress neuropeptide family, and its receptor CRF type 2 (CRFR2) signalling in the MePD are involved in mediating the suppressive effect of psychosocial stress on LH pulsatility (3). However, the underlying neural mechanisms involved in mediating the inhibitory effect of MePD Ucn3 neurons on LH pulsatility remain to be established. GABA and glutamate signalling in the amygdala are implicated in the modulation of anxiety behaviour (15,16). The amygdala also modulates the stress response through the hypothalamic pituitary adrenal (HPA) axis. The medial amygdala activates the HPA axis in response to predator odor stress (17) and the MePD in particular has been shown to send stress-activated efferents to the paraventricular nucleus of hypothalamus (PVN) (18). Furthermore, MePD Ucn3 and CRFR2-positive neurons are implicated in regulating the stress response, with restraint stress increasing MePD Ucn3 (19) and social defeat elevating MePD CRFR2 expression in rodents (20). Moreover, MePD Ucn3 and CRFR2-positive neurons project to the PVN (21), and we have recently shown that chemogenetic inhibition of MePD Ucn3 neurons prevents psychogenic stress-induced corticosterone (CORT) release in female mice (3).

In this study, we aimed to determine whether activation of MePD Ucn3 neurons inhibits pulsatile LH secretion via GABA and or glutamate signalling within this amygdala subnucleus. To achieve this, we will combine selective optogenetic activation of MePD Ucn3 neurons with pharmacological antagonism of GABA_A_ or GABA_B_ receptors, or a combination of N-methyl-D-aspartate (NMDA) and α-amino-3-hydroxy-5-methyl-4-isoxazole propionic acid (AMPA) receptors, respectively, using chronically implanted intra-MePD optofluid cannulae while collecting serial blood samples for LH pulse measurement in Ucn3-Cre-tdTomato female mice. Additionally, we will investigate whether optogenetic stimulation of MePD Ucn3 projection terminals in the PVN induce CORT release and suppress LH pulsatility.

## Methods

### Mice

Cryopreserved sperm of Ucn3-cre mice (strain Tg(Ucn3-cre)KF43Gsat/Mmucd; congenic on C57BL/6 background) was acquired from MMRRC GENSAT and heterozygous transgenic breeding pairs of Ucn3-Cre mice were recovered via insemination of female C57Bl6/J mice at King’s College London, as described previously (3). Ucn3-Cre mice were genotyped using PCR for the detection of heterozygosity and Ucn3-cre mice were bred with td-Tomato mice (strain B6.Cg-Gt(ROSA)26Sortm9(CAG-tdTomato)Hze/J; congenic on C57BL/6 background) acquired from The Jackson Laboratory, Bar Harbor, ME, USA to produce Ucn3-cre-tdTomato reporter mice, as described previously (3,21). Female Ucn3-cre-tdTomato mice weighing 19-23 g and aged 6-8 weeks were singly housed in individually ventilated cages sealed with a HEPA-filter at 25 ± 1 °C in a 12:12 h light/dark cycle, lights on at 07:00 h equipped with nesting material, wood-chip bedding and food and water ad libitum. All procedures were carried out following the United Kingdom Home Office Regulations and approved by the Animal Welfare and Ethical Review Body Committee at King’s College London.

### Stereotaxic injection of adeno-associated-virus carrying the channelrhodopsin construct and fibre optic or optofluid cannula implantation

All surgical procedures were carried out under aseptic conditions with general anaesthesia using ketamine (Vetalar, 100 mg/kg, i.p.; Pfizer, Sandwich, UK) and xylazine (Rompun, 10 mg/kg, i.p.; Bayer, Leverkusen, Germany). Ucn3-Cre-tdTomato mice were secured in a David Kopf stereotaxic frame (Kopf Instruments, Model 900) and either solely bilaterally ovariectomised (OVX) or implanted with an 17β-estradiol (E_2_) silastic capsule. E_2_ was dissolved in sesame oil to reach a final concentration of 36 μg E_2_/mL and filled in 14 mm capsules of inner/outer diameter: 1.575/3.175 mm, as previously described (22). The reference study reported that at 21 days of implantation, serum E_2_ concentrations were within physiological concentrations equivalent to those found in mice in the diestrous stage of the cycle. However, by 35 days of implantation the serum E_2_ concentrations had slowly declined but still within the range found in diestrous mice. Due to the technically demanding nature of our experiments the E_2_ capsules were implanted for longer than 35 days post-surgery during the experimental period. Therefore, E_2_ levels may be lower than expected for the diestrous phase. E_2_ is known to enhance the suppressive effects of stress where the action of the gonadal steroids sensitizes the HPA axis and augments stress-induced inhibition of the GnRH pulse generator in many species, from rodents to primates (42–45). We had two experimental groups; OVX only and OVX + E_2_ capsule. The mouse brain atlas of Paxinos and Franklin (23) was used to obtain target coordinates for the MePD (2.30 mm lateral, −1.55 mm from bregma, at a depth of −4.94 mm below the skull surface). To reveal the skull an incision was made of the scalp and one small hole was drilled above the location of the right MePD. AAV9-EF1a-double floxed-hChR2(H134R)-EYFP-WPRE-HGHpA (300 nL; 3×10^11^ GC/ml; Serotype:9; Addgene, Massachusetts, USA) was unilaterally injected into the right MePD using a 2-μL Hamilton micro syringe (Esslab, Essex, UK) over 10 min, using the robot stereotaxic system (Neurostar, Tubingen, Germany), performed for the targeted expression of ChR2-EYFP in MePD Ucn3 neurons. After injection, the needle was left in position for a further 5 min and lifted slowly over 2 min. Cre-positive mice received the AAV-ChR2 injection (test mice) or a control virus AAV-YFP (Addgene) (control mice). The control virus does not contain the ChR2 construct. The mice were then implanted with a dual optofluid cannula (Doric Lenses, Quebec, Canada) at the same AP and ML coordinates as the viral injection site, however a different DV was used such that the internal cannula targets the MePD and the fibre optic cannula is situated 0.2 mm above the MePD site. Once in position the optofluid cannula was secured on the skull using dental cement (Super-Bond Universal Kit, Prestige Dental, UK) and the incision of the skin was closed with suture. Mice were left to recover for one week and after recovery period, mice were handled daily to acclimatise to experimental procedures for a further 2 weeks. Thirteen mice received the AAV-ChR2 injection and were implanted with an optofluid cannula in the MePD. Five mice received the control AAV-YFP and were implanted with an optofluid cannula in the MePD. Eleven mice received the AAV-ChR2 injection in the MePD and were implanted with an optic fibre in the PVN. Five mice received the control AAV-YFP and were implanted with an optic fibre in the PVN.

### In vivo optogenetic stimulation of MePD Urocortin-3 neurons

In order to test the effect of optogenetic stimulation of MePD Ucn3 neurons on LH pulsatility the ferrule of the implanted optofluid cannula, on the right side of the MePD, was attached via a ceramic mating sleeve to a multimode fibre optic rotatory joint patch cable (Thorlabs LTD, Ely, UK) at a length allowing for free movement of the mouse in their cage and blue light (473 nm wavelength; 10 mW) was delivered using a Grass SD9B stimulator-controlled laser (Laserglow Technologies, Toronto, Canada). The mice were left to acclimatise for 1 h. Following the acclimatisation period blood samples (5 μL) were collected every 5 min for 2 h where after 1 h of control blood sampling, Ucn3-cre-tdTomato mice received optic stimulation at 10 Hz with a 10 ms pulse interval and pattern stimulation of 5 seconds on and 5 seconds off for the remaining 1 h.

### Intra-MePD administration of bicuculline (BIC), CGP-35348 or AP5 + CNQX during optogenetic stimulation of MePD Urocortin-3 neurons

Neuropharmacological manipulation of GABA and glutamate receptor signalling in the MePD with or without optogenetic stimulation of MePD Ucn3 neurons was done to test whether MePD Ucn3 induced suppression of LH pulsatility involves GABA or glutamate receptor signalling. OVX mice implanted with a optofluid cannula in the MePD were subjected to the tail-tip blood collection procedure, as previously described (24). Infusion of drugs with or without optogenetic stimulation and blood sampling were performed between 09:00-13:00 h, where 5 μl blood was collected every 5 min for 2 h. Internal cannula (Doric Lenses) attached to extension tubing (0.58 mm ID, 0.96 mm OD), preloaded with bicuculline (BIC) (GABA_A_ receptor antagonist; Sigma-Aldrich), CGP-35348 (GABA_B_ receptor antagonist; Sigma-Aldrich), AP5 + CNQX cocktail (NMDA and AMPA receptor antagonists; Tocris) or aCSF as vehicle control, was inserted into the guide cannula of the optofluid implant and mice were connected to the laser as described above. The tubing for drug infusion extended beyond the cage and the distal ends were attached to 10-μl Hamilton syringes (Waters Ltd, Elstress, UK) fitted into a PHD 2000 Programmable syringe pump (Harvard Apparatus, Massachusetts, USA), allowing for a continuous infusion of the drug at a constant rate and mice were kept in the cage throughout the experiment, freely-moving with food and water *ad libitum*. After 55 min of control blood sampling, the mice were given an initial bolus injection, 0.30 μl at a rate of 0.06 μl/min over 5 min, of BIC, CGP-35348, AP5 + CNQX or aCSF. The laser was turned on at 60 min and the bolus injection was followed by a continuous infusion, 0.8 μl at a rate of 0.01 μl/min over 1 h, of BIC, CGP-35348, AP5 + CNQX or aCSF. BIC, CGP-35348, AP5 and CNQX were dissolved in aCSF to reach bolus concentrations of 20 pmol, 4.5 nmol, 1.2 nmol and 0.5 nmol respectively. The concentrations for the continuous infusion of BIC, CGP-35348, AP5 and CNQX were 68 pmol, 15 nmol, 2nmol and 1 nmol respectively. In the absence of optic stimulation, the same regimen was applied. Data collection was conducted over a period of 4-6 weeks. The mice receive all treatments in a random order, with at least 2 days between experiments.

### MePD Urocortin-3 terminal optogenetic stimulation in the PVN

To test the effect of optogenetic stimulation of MePD Ucn3 neuronal terminals in the PVN on LH pulsatility and CORT release, OVX + E_2_ Ucn3-cre-tdTomato mice with AAV-ChR2 injection in the right MePD implanted with a fibre optic cannula in the right PVN were subjected to the tail-tip blood collection procedure, as previously described (24). For LH measurement, blood samples were collected every 5 min for 2 h, as described above. Mice were connected to the laser as described above. Optogenetic stimulation was initiated after the 1 h control blood sampling period and Ucn3-cre-tdTomato mice received optic stimulation at 20 Hz, 15 mW with a 10 ms pulse interval and pattern stimulation of 5 seconds on and 5 seconds off for the remaining 1 h. For CORT measurements, 15 μl blood samples were collected on a separate occasion and blood samples were stored in tubes containing 5 μl heparinised saline (5 IU ml^-1^). Optogenetic stimulation was initiated, as described above, 30 min into the experiment (time point, 0 min) and lasted for 1 h (time point 60 min). Remaining blood samples were collected at 0, 30 and 60 min, with a final sample taken 1 h after the termination of optogenetic stimulation (time point, 120 min). At the end of the experiment, blood samples were centrifuged at 13,000 RPM for 20 min at 20°C and plasma stored at −20°C. Data collection was conducted over a period of 4-6 weeks. All experiments were performed between 9:00-12:00 h.

### Validation of AAV injection and cannula implant site

Upon completion of experiments, mice were anaesthetised with a lethal dose of ketamine and transcardial perfusion was performed with heparinised saline for 5 min followed by ice-cold 4% paraformaldehyde (PFA) in phosphate buffer (pH 7.4) for 15 min using a pump (Minipuls, Gilson, Villiers Le Bel, France). Mice brains were collected immediately and post-fixed in 15% sucrose in 4% PFA at 4 °C and left to sink. Once sunk, they were transferred to 30% sucrose in phosphate-buffered saline (PBS) and left to sink. Brains were snap-frozen in isopropanol on dry ice and stored in −80°C until further processing. Every third coronal brain section (30-μm/section) through-out the MePD region (−1.34 mm to −2.70 mm from bregma) was collected using a cryostat (Bright Instrument Co., Luton, UK). Sections were mounted on microscope slides, air dried and covered with ProLong Antifade mounting medium (Molecular Probes, Inc. OR, USA). Cannula placement and AAV injection site was verified with Axioskop 2 Plus microscope equipped with Axiovision, version 4.7 (Zeiss) by determining whether cannula reach the MePD. For AAV-ChR2-EYFP injected Ucn3-cre-tdTomato mice we determined whether Ucn3 neurons were infected in the MePD region by merging td-Tomato fluorescence of Ucn3 neurons with YFP fluorescence in the MePD. The number of Ucn3 mCitrine-positive neurones co-localised with td-Tomato fluorescence in the MePD of each animal was counted using 4 sections and the average number of neurons presented is per section per MePD. The group mean percent was calculated by taking the average number of Ucn3 mCitrine-positive neurons out of the average number of Ucn3 neurons expressing td-Tomato fluorescence per 4 sections and presented as mean ± SEM %, as described previously (3). Axioskop 2 Plus microscope (Carl Zeiss) equipped with axiovision, version 4.7 (Carl Zeiss) was used to take images. Only mice with correct AAV injection and cannula placement in the MePD were analysed.

### LH and CORT measurement

Mice were handled daily to acclimatise to the tail-tip blood sampling procedure for LH measurement (3). Blood samples collected for LH measurement were processed using LH ELISA, as previously reported (3). The capture antibody (monoclonal antibody, anti-bovine LHβ subunit, <u>AB_2665514</u>) was purchased from Department of Animal Science at the University of California, Davis. The mouse LH standard (AFP-5306A) and primary antibody (polyclonal antibody, rabbit LH antiserum, <u>AB_2665533</u>) were obtained from Harbour-UCLA (California, USA). The secondary antibody (Horseradish-Peroxidase (HRP)-linked donkey anti-rabbit IgG polyclonal antibody, <u>AB_772206</u>) was purchased from VWR International (Leicestershire, UK). Intra-assay and inter-assay variations were 4.6% and 10.2%, respectively and the assay sensitivity was 0.0015 ng/mL. DynPeak algorithm was used for the detection of LH pulses (3). The effect of optogenetic stimulation of MePD Ucn3 neurons and neuropharmacology studies was established by comparing the LH inter-pulse interval (IPI) from the 1 h control period (pre-stimulation) or drug administration control period to the 1 h experimental period, as previously described (3). The LH pulse amplitude was calculated as the difference between the peak of an LH pulse and the baseline LH level before the onset of the pulse. The means were compared between the pre-treatment and treatment periods for all experiments. The LH pulse amplitude was also compared between experimental groups within the treatment period. The mean LH value for each LH pulse profile was quantified by summing all values from one LH pulse profile and dividing by the number of values recorded in 120 mins.

Blood samples were collected for CORT measurement were processed using a commercially available enzyme immunoassay (EIA) kit (sheep polyclonal antibody specific for corticosterone, <u>AB_2877626</u>; DetectX^®^ Corticosterone Enzyme Immunoassay Kit, K014; Arbor Assays, Michigan, USA), as described previously (3).

### Statistics

Mice were compared between groups using a two-way ANOVA with a Tukey post-hoc and data was presented as mean ± SEM. Statistics were performed using Igor Pro 7, Wavemetrics, Lake Oswego, OR, USA. Data was represented as mean ± SEM and +p<0.05, ++p<0.001 and +++p<0.0001 were considered to be significant.

## Results

### Effects of optogenetic stimulation of MePD Urocortin-3 neurons on LH pulse frequency

Optogenetic stimulation of MePD Ucn3 neurons in control mice had no effect on LH pulse interval (Fig. 1A; AAV-YFP: n = 3) and administration of aCSF alone (AAV-YFP with no stimulation: n = 2) had no effect on LH pulse interval. Optogenetic stimulation of MePD Ucn3 neurons with blue light at 10 Hz, 10 mW suppressed pulsatile LH secretion (Fig. 1B; AAV-ChR2: n = 9). These data are summarized in the figure 1C. LH pulse amplitude was quantified and data is provided in supplemental table (25).

**Figure 1.**
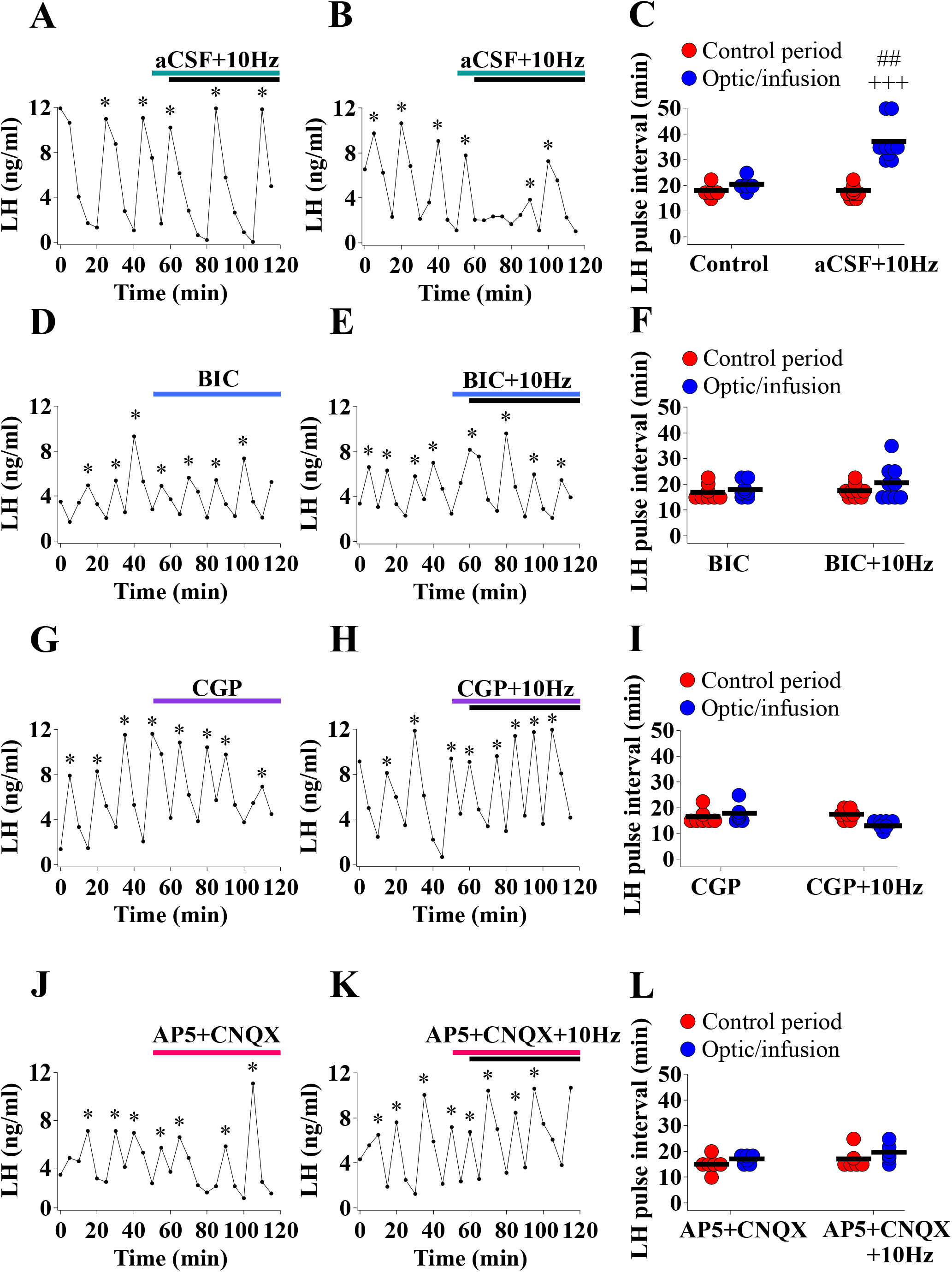
Optogenetic stimulation of posterodorsal medial amygdala (MePD) urocortin-3 (Ucn3) neurons with blue light at 10 Hz, 10 mW suppressed pulsatile LH secretion in adult ovariectomised (OVX) Ucn3-cre-tdTomato female mice. Infusion of bicuculline (BIC), GABAA receptor antagonist, CGP, GABAB receptor antagonist, and AP5 and CNQX, NMDA and AMPA receptor antagonists, respectively, into the posterodorsal medial amygdala (MePD) during optogenetic stimulation of MePD urocortin-3 (Ucn3) neurons with blue light at 10 Hz, 10 mW completely blocked the suppressive effect of MePD Ucn3 neuronal activation on LH pulsatility. Representative LH pulse profiles showing the effects of (A) control AAV-YFP mice administered with aCSF receiving optical stimulation of 10 Hz, 10 mW, (B) AAV-ChR2 mice administered with aCSF receiving optical stimulation of 10 Hz, 10 mW, (C) Summary of LH pulse interval for the pre-stimulation control period (1 h) and optical stimulation/infusion period (1 h), (D) AAV-ChR2 mice administered with BIC, (E) AAV-ChR2 mice administered with BIC receiving optical stimulation of 10 Hz, 10 mW, (F) Summary of LH pulse interval, (G) AAV-ChR2 mice administered with CGP, (H) AAV-ChR2 mice administered with CGP receiving optical stimulation of 10 Hz, 10 mW, (I) Summary of LH pulse interval, (J) AAV-ChR2 mice administered with a combination of AP5 and CNQX, (K) AAV-ChR2 mice administered with a combination of AP5 and CNQX receiving optical stimulation of 10 Hz, 10 mW, (L) Summary of LH pulse interval. LH pulses detected by the DynePeak algorithm are indicated with an asterisk located above each pulse on the representative LH pulse profiles. For A-C +++p<0.0001 control period vs optic stimulation + aCSF (AAV-ChR2: n = 9); ##p<0.001 optic stimulation vs control group (AAV-YFP: n = 3; no stimulation: n = 2). For D-F n = 8-9 per group. For G-I n = 7 per group. For J-L n = 6 per group.

### Effects of intra-MePD administration of bicuculline (BIC), a GABAAR antagonist, and CGP, a GABA_B_R antagonist, during optical stimulation of MePD Urocortin-3 neurons on LH pulsatility

Delivery of BIC alone had no significant effect on LH pulsatility (Fig. 1D; n = 8). Intra-MePD infusion of BIC during optogenetic stimulation of MePD Ucn3 neurons with blue light at 10 Hz, 10 mW completely blocked the suppressive effect of MePD Ucn3 neuronal activation on LH pulsatility (Fig. 1E; n = 9). The results of this experiment are summarised in the figure 1F. LH pulse amplitude was quantified and data is provided in supplemental table (25).

### Effects of intra-MePD administration of CGP, a GABABR antagonist, during optical stimulation of MePD Urocortin-3 neurons on LH pulsatility

Delivery of CGP alone had no significant effect on LH pulsatility (Fig. 1G; n = 7). Intra-MePD infusion of CGP during optogenetic stimulation of MePD Ucn3 neurons with blue light at 10 Hz, 10 mW completely blocked the suppressive effect of MePD Ucn3 neuronal activation on LH pulsatility (Fig. 1H; n = 7). The results of this experiment are summarised in the figure 1I. LH pulse amplitude was quantified and data is provided in supplemental table (25).

### Effects of intra-MePD glutamate antagonism during optogenetic stimulation of MePD Urocortin-3 neurons on LH pulsatility

Delivery of AP5 + CNQX alone had no significant effect on LH pulsatility (Fig. 1J; n = 6). Intra-MePD infusion of AP5 (selective NMDA antagonist) + CNQX (selective AMPA antagonist) during optogenetic stimulation of MePD Ucn3 neurons with blue light at 10 Hz, 10 mW completely blocked the suppressive effect of MePD Ucn3 neuronal activation on LH pulsatility (Fig. 1K; n = 6). The results of this experiment are summarised in the figure 1L. LH pulse amplitude was quantified and data is provided in supplemental table (25).

### Effect of MePD Urocortin-3 terminal stimulation in the PVN on dynamic CORT secretion

Optogenetic stimulation of MePD Ucn3 neuron terminals in the PVN with blue light at 20 Hz, 15 mW elevated CORT secretion compared to controls (Fig. 2; AAV-ChR2: n = 6). Optogenetic stimulation of MePD Ucn3 neurons and terminal projections in the PVN in control mice had no effect on dynamic CORT secretion (AAV-YFP; n = 5).

**Figure 2.**
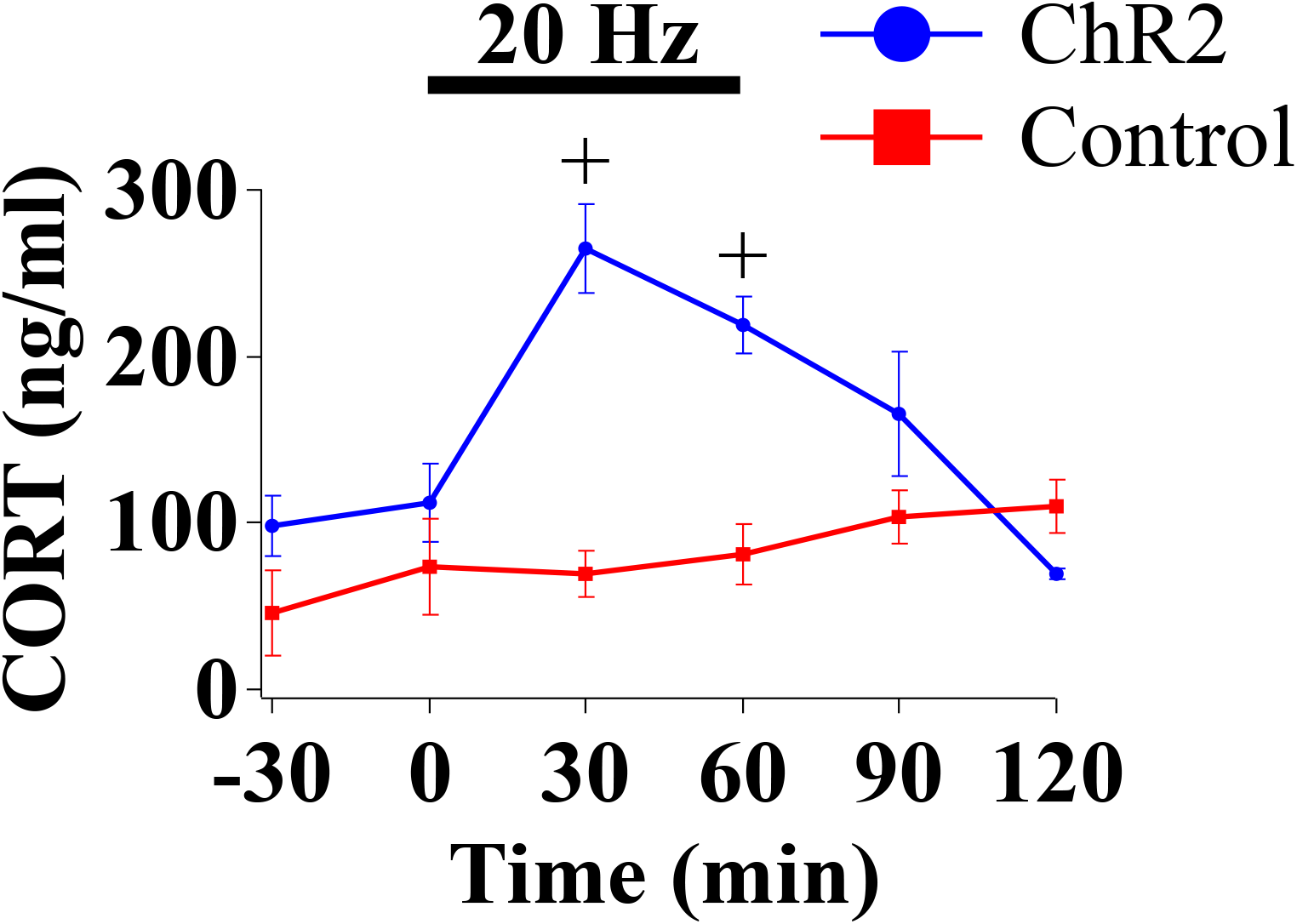
Optogenetic stimulation of posterodorsal medial amygdala (MePD) urocortin-3 (Ucn3) neuronal terminals in the hypothalamic paraventricular nucleus (PVN) with blue light at 20 Hz, 15 mW elevated corticosterone (CORT) secretion in adult ovariectomised (OVX) + 17β-estradiol (E_2_) Ucn3-cre-tdTomato female mice. CORT secretion time-course for mice receiving optical stimulation initiated at 0 min and terminated at 60 min (1 h duration), followed by a 1 h recovery period (60-120 min) for the AAV-ChR2 group (blue line, circles) and control group (red line, squares). +p<0.05 AAV-ChR2 (n = 6) at time point 30 min and 60 min vs. control (AAV-YFP: n = 5) at time point 30 min and 60 min.

### Effect of MePD Urocortin-3 terminal stimulation in the PVN on LH pulsatility

Optogenetic stimulation of MePD Ucn3 projections in the PVN in control mice had no effect on LH pulse interval (Fig. 3A and C; AAV-YFP, n = 3; wild-type with AAV-ChR2, n = 2) and no stimulation of the MePD Ucn3 projections in the PVN (no stimulation, n = 2) had no effect on LH pulse interval. Optogenetic stimulation of MePD Ucn3 neuron terminals in the PVN with blue light at 20 Hz, 15 mW reduced LH pulsatility compared to controls (Fig. 3B and D; AAV-ChR2: n = 8). These data are summarised in the figure 3E. LH pulse amplitude was quantified and data is provided in supplemental table (25).

**Figure 3.**
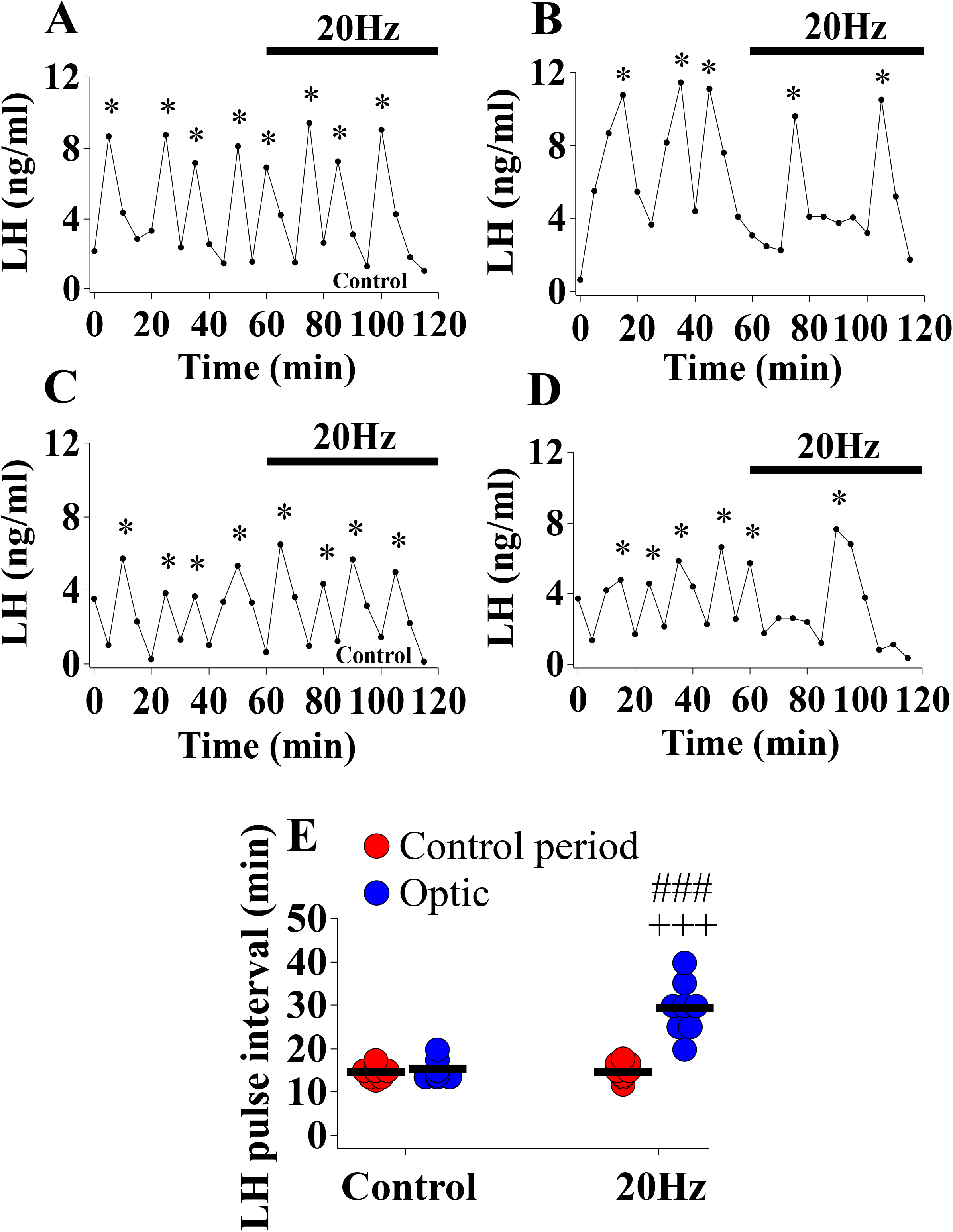
Optogenetic stimulation of posterodorsal medial amygdala (MePD) urocortin-3 (Ucn3) neuronal terminals in the hypothalamic paraventricular nucleus (PVN) with blue light at 20 Hz, 15 mW reduced LH pulsatility in adult ovariectomised (OVX) + 17β-estradiol (E_2_) Ucn3-cre-tdTomato female mice. Representative LH pulse profiles showing the effects of (A) AAV-YFP/WT mice receiving optical stimulation of 20 Hz, 5 mW, (B) AAV-ChR2 mice receiving optical stimulation of 20 Hz, 5 mW, (C) Another example of AAV-YFP/WT mice receiving optical stimulation of 20 Hz, 5 mW, (D) Another example of AAV-ChR2 mice receiving optical stimulation of 20 Hz, 5 mW, (E) Summary of LH pulse interval for the pre-stimulation control period (1 h) and optic stimulation period (1 h). LH pulses detected by the DynePeak algorithm are indicated with an asterisk located above each pulse on the representative LH pulse profiles. +++p<0.0001 control period vs optic stimulation for AAV-ChR2 mice (n = 8); ###p<0.0001 AAV-YFP/WT (AAV-YFP, n = 3; wild-type with AAV-ChR2, n = 2; no stimulation, n = 2) vs AAV-ChR2 mice during optic stimulation.

### Selective expression of ChR2 in MePD Ucn3 neurons and MePD Ucn3 neuron projections in the PVN, and validation of cannula position

Evaluation of AAV9-EF1a-double floxed-hChR2(H134R)-EYFP-WPRE-HGHpA expression in tdTomato labelled neurons from AAV-injected Ucn3-cre-tdTomato mice revealed that 87.00 ± 6.00% of MePD Ucn3 neurons expressed AAV-ChR2 and the number of tdTomato labelled Ucn3 neurons per side per slice in the MePD was counted at 49.00 ± 5.37 (mean ± SEM) with the number of AAV-ChR2 infected neurons being 43.01 ± 3.65 (mean ± SEM) (n=8). Non-specific expression of ChR2 in the MePD (YFP single labelling) has been quantified to be limited to 6 ± 4.63 (mean ± SEM) in all of 8 mice. A representative example is shown in figure 4, A-F. The optofluid cannula was also verified to be correctly positioned in the MePD in all of 8 mice. Evaluation of AAV9-EF1a-double floxed-hChR2(H134R)-EYFP-WPRE-HGHpA expression in Ucn3 projections from the MePD to the PVN showed YFP labelled fibres in the PVN figure 5, A-Aiii. Local Ucn3 neurons in the PVN are labelled with tdTomato. We cannot however exclude the possibility that a small portion of fibres may be present that are not colocalized with tdTomato, which may reflect unspecific expression of ChR2 in nerve terminals other than Ucn3 in the PVN. The optic fibre cannula was also verified to be correctly positioned in the PVN in all of 8 mice.

**Figure 4.**
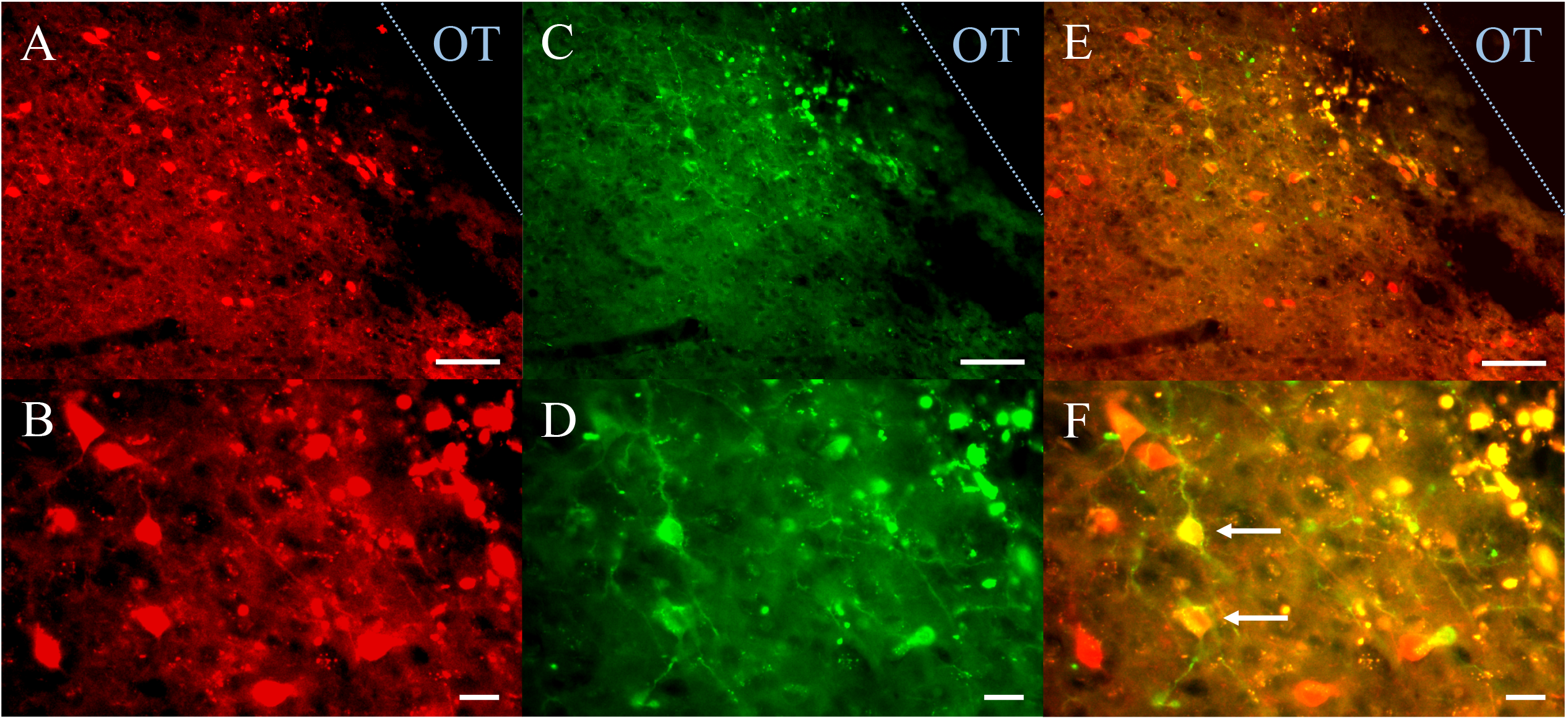
Expression of AAV9-EF1a-double floxed-hChR2(H134R)-EYFP-WPRE-HGHpA in posterodorsal medial amygdala (MePD) Urocortin-3 (Ucn3) neurons. (A-E) Representative dual fluorescence photomicrographs of the MePD from a Ucn3-cre-tdTomato female ovariectomised (OVX) mouse injected with AAV9-EF1a-double floxed-hChR2(H134R)-EYFP-WPRE-HGHpA. Ucn3 neurons labelled with tdTomato (A) and YFP (C) appear yellow/orange (E). (B), (D) and (F) are a higher power view of (A), (C) and (E) respectively. Scale bars represent A, C, E 50 μm and B, D, F 20 μm. White arrows show examples of Ucn3 neurons labelled with tdTomato and YFP. OT, Optic tract (blue line).

**Figure 5.**
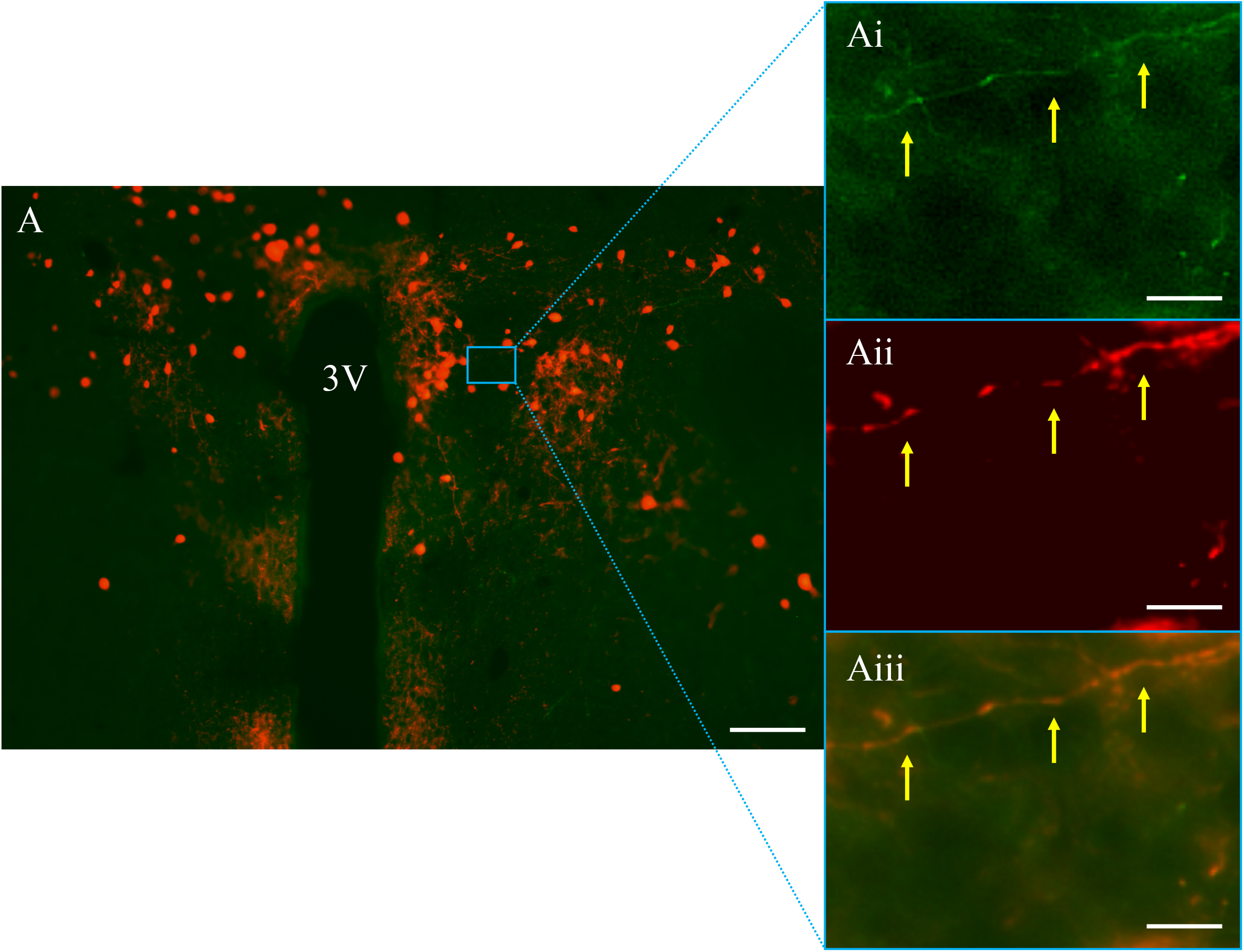
Expression of AAV9-EF1a-double floxed-hChR2(H134R)-EYFP-WPRE-HGHpA in Urocortin-3 (Ucn3) neuronal projection in the paraventricular nucleus of the hypothalamus (PVN) originating from the posterodorsal medial amygdala (MePD). (A-Aiii) Representative dual fluorescence photomicrographs of the PVN from a Ucn3-cre-tdTomato female ovariectomised (OVX) + 17β-estradiol (E_2_) mouse injected with AAV9-EF1a-double floxed-hChR2(H134R)-EYFP-WPRE-HGHpA. Ucn3 neurons and projections labelled with (A) YFP and tdTomato expression in PVN, (Ai) higher power view of YFP labelled MePD Ucn3 projections in the PVN, (Aii) higher power view of tdTomato labelled MePD Ucn3 projections in the PVN and (Aiii) YFP and tdTomato merged. Scale bars represent A, 100 μm, Ai, Aii and Aiii, 10 μm. Blue box shows the YFP labelled MePD Ucn3 projections in the PVN, yellow arrows point to parts of the YFP labelled MePD Ucn3 projection in the PVN. 3V; third ventricle.

## Discussion

We show optogenetic stimulation of MePD Ucn3 neurons inhibits pulsatile LH secretion, an effect that was completely blocked by intra-MePD GABA and glutamate antagonism. Moreover, optogenetic stimulation of MePD Ucn3 projections in the hypothalamic PVN also suppresses LH pulsatility while inducing the release of CORT. These results demonstrate that inhibition of the GnRH pulse generator by Ucn3 neurons within the MePD relies on GABA and glutamate signalling in the MePD, *per se*. Moreover, MePD Ucn3 efferent projections in the PVN modulate the activity of the HPA and HPG axes.

The MePD is a major upstream regulator of gonadotropic hormone secretion with a significant population of GABAergic neurons projecting to reproductive neural centres in the hypothalamus (26). Although classic lesioning studies suggest an inhibitory output of the MePD over reproductive function (27,28), selective optical stimulation of MePD kiss1 neurons increases LH pulse frequency (4), which our unpublished observations show relies upon both GABA and glutamate signalling in the MePD. Due to this interesting dichotomy, we have proposed a disinhibitory system, which is consistent with the pallidal origin of this limbic structure (29), whereby MePD kiss1 activation excites inhibitory GABAergic interneurons, which in turn reduces the inhibitory tone of the GABAergic projections from the MePD to the ARC KNDy network allowing for an increase in GnRH pulse generator frequency (4). Contrastingly, kiss1R antagonism, reducing MePD kiss1 signalling decreases GnRH pulse generator frequency (30).

We have recently shown that stress-induced activation of Ucn3 and CRFR2 signalling within the MePD inhibits GnRH pulse generator activity (3) and activation of MePD Ucn3 neurons delays pubertal timing (31). It has also been shown that the majority of MePD CRFR2 expressing neurons are GABAergic (21). In the MePD, Ucn3 fibres overlap with sites expressing its cognate receptor CRFR2 (32,33) and some local Ucn3 neurons within the MePD are shown to have an interneuron-like appearance (34), thus Ucn3 neurons may form connections with and signal via GABAergic CRFR2-positive neurons in this region. In the present study, we show that optogenetic activation of MePD Ucn3 neurons inhibits pulsatile LH secretion. Therefore, we propose that activated MePD Ucn3 neurons may signal through inhibitory GABAergic CRFR2-positive interneurons, upstream to the MePD kiss1 neurons, downregulating MePD kiss1 signalling resulting in decreased GnRH pulse generator frequency, as illustrated in figure 6A.

**Figure 6.**
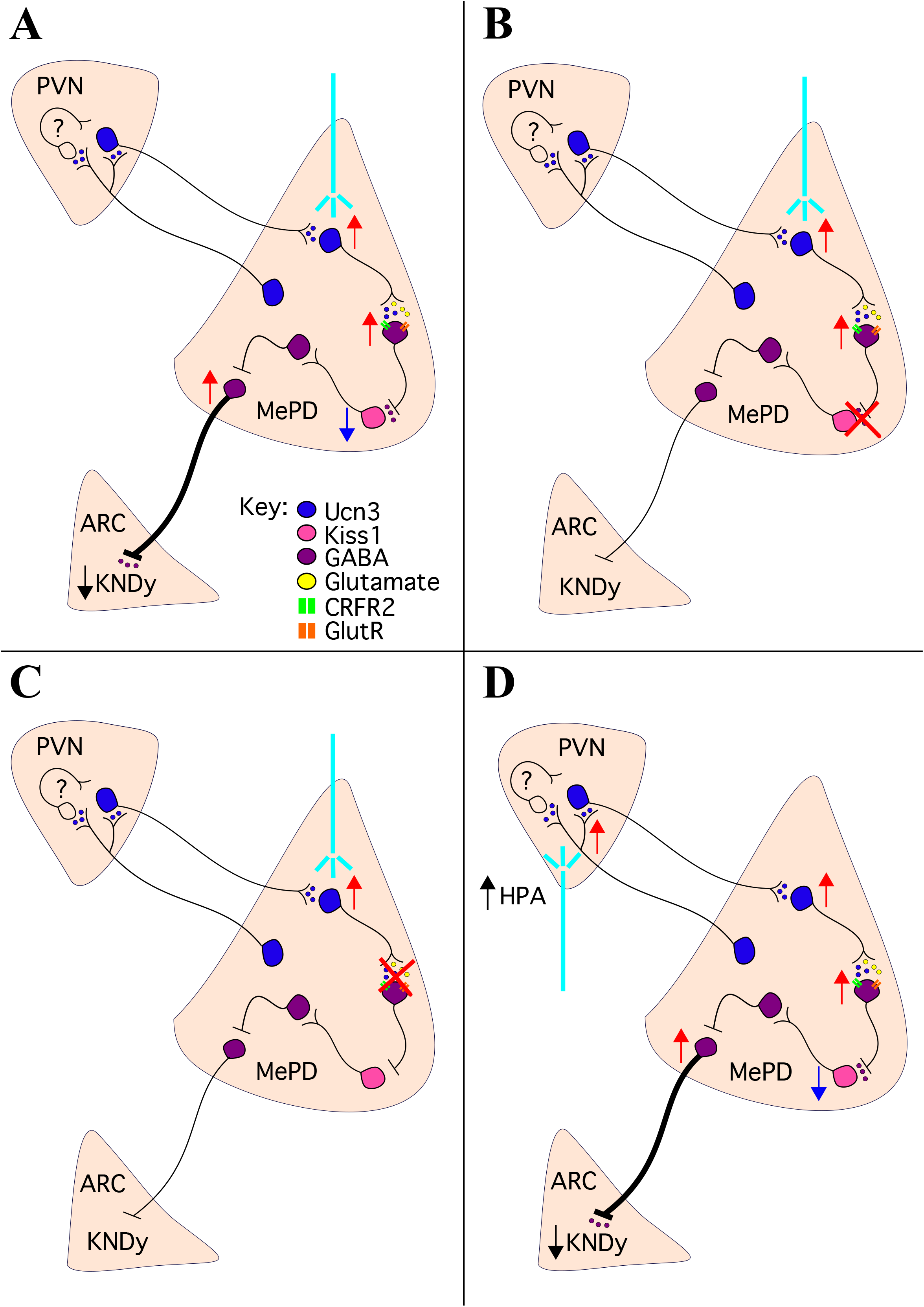
Schematic illustration of the MePD neurocircuits and their pharmacological manipulation. (A) Optical stimulation of local MePD Ucn3 neurons releases Ucn3 and the co-transmitter glutamate, which activate inhibitory GABAergic CRFR2-positive interneurons located upstream to the kiss1 neurons suppressing their activity. The reduced kiss1 signalling deactivates its downstream inhibitory GABAergic interneuron causing disinhibition of the GABAergic projection efferents from the MePD, resulting in an increased inhibitory tone over the ARC KNDy network, thus decreasing GnRH pulse generator frequency. (B) Antagonism of both GABA_A_ and GABA_B_ receptors in the MePD during optical stimulation of Ucn3 neurons blocks the downregulation of kiss1 neuronal activity, preventing the inhibitory effect of Ucn3 signalling on the KNDy network. (C) Antagonism of both NMDA and AMPA receptors in the MePD during optical stimulation of Ucn3 neurons blocks the activation of the downstream GABAergic CRFR2-positive interneurons, preventing the inhibitory effect of Ucn3 signalling on the KNDy network and suggesting glutamate is necessary for the inhibition of pulsatile LH secretion. (D) High frequency photoactivation of MePD Ucn3 projection terminals in the PVN may activate Ucn3 neurons and/or other neurons of an unknown phenotype in the PVN resulting in stimulation of the HPA axis and CORT release. Additionally, there may be collateral activation of reciprocal PVN Ucn3 projections to the MePD, which in turn relay via the MePD Ucn3 neuronal circuitry an inhibitory influence on the ARC KNDy neuronal network to suppress GnRH pulse generator frequency. Urocortin3 (Ucn3), Corticotropin-releasing factor type 2 receptor (CRFR2), glutamate receptor (GlutR), the posterodorsal subnucleus of the medial amygdala (MePD), paraventricular nucleus of hypothalamus (PVN), the hypothalamic arcuate nucleus (ARC).

In our proposed model, we hypothesised that GABA signalling is an important factor mediating the suppressive effect of MePD Ucn3 activation on GnRH pulse generator activity. Therefore, we tested the effect of intra-MePD administration of GABAA or GABAB receptor antagonists during optical stimulation of MePD Ucn3 neurons on LH pulsatility, and indeed, we found that both GABA_A_ and GABA_B_ receptor antagonism blocks the suppression of LH pulses by MePD Ucn3 neuronal activation. Indeed, it has been shown that in GABAB receptor knock out mice Kiss1 mRNA expression increases in the MePD suggesting GABA_B_ regulates Kiss1 expression in this region (35). Whether the MePD kiss1 neurons also contain GABA_A_ receptors is unknown. Selective optical stimulation of Ucn3 neurons may activate inhibitory GABAergic CRFR2-positive interneurons (35), which in turn downregulate MePD kiss1 signalling, which we have previously shown results in suppression of GnRH pulse generator frequency (30) (see Fig. 6A). Although, based on our proposed neuronal circuit, GABA_A_ and GABA_B_ receptor antagonism may be working upstream and downstream of MePD kiss1 activity to cancel both the inhibitory GABAergic CRFR2 interneurons upstream to and the inhibitory GABAergic interneurons downstream to MePD kiss1 neurons. Therefore, the inhibitory output of the MePD GABAergic projection neurons to the ARC KNDy network is cancelled, thus, negating the suppressive effect of MePD Ucn3 activation on the GnRH pulse generator. Essentially, terminating the ability of the GABA interneuron, upstream and downstream of MePD kiss1 activity, to take part in the disinhibition by antagonising GABA receptors located on the MePD kiss1 neurons as well as post-synaptic GABA projection neurons removing the inhibitory control over the GnRH pulse generator from the MePD (see Fig. 6B).

We have identified GABA signalling as a key part of the MePD Ucn3 neural circuitry associated with suppression of the GnRH pulse generator, however glutamate signalling in the amygdala is also known to be involved in modulating reproduction and anxiety-like behaviour (36). Administration of NMDA into the MePD disrupts ovarian cycles in rats, while NMDA receptor antagonism in the MePD delays pubertal timing (28). We hypothesised that glutamate neurotransmission may form a critical part of the MePD Ucn3 neural circuitry involved in stress-induced inhibition of GnRH pulse generator activity. In the present study, we show that intra-MePD AMPA and NMDA receptor antagonism blocks the inhibitory effect of MePD Ucn3 neuronal activation on LH pulsatility. These observations suggest the involvement of similar neural circuitry as in the neighbouring posteroventral medial amygdala which exhibits functional glutamatergic signalling onto inhibitory GABA neurons (29). Moreover, inhibition of both GABA and glutamate signalling is equally effective at blocking suppression of LH pulses suggesting they are key components in the circuit.

We know the MePD Ucn3 neurons are interconnected with each other as well as CRFR2 expressing neurons, forming a Ucn3-CRFR2 neuronal circuit (21). The effect of glutamate antagonism illustrated in figure 6C, suggests that glutamate transmission may be mediated via glutamatergic Ucn3 neurons. This has been shown in the hypothalamic perifornical area, where they are activated during infant-directed aggression (37) and increase anxiety-like behaviour (38), but this awaits future study for the MePD Ucn3 population. Our previous data shows that infusing Ucn3 in the MePD blocks LH pulses indicating their activity in this brain region is sufficient to suppress GnRH pulse generator activity (3). We have also shown that infusion of CRFR2 antagonists in the MePD blocks the stress-induced suppression of LH pulses suggesting MePD CRFR2 expressing neurons are also involved in modulating the GnRH pulse generator under stressful conditions. Our current data expands on our previous observations determining whether neurotransmission modulating pulsatile LH secretion is involved in this MePD circuit. Our results suggest that glutamate may be necessary for the inhibition of LH pulses induced by activation of MePD Ucn3 neurons, thus we show that glutamate signalling is an indispensable part of this circuit. Based on our observations, we propose that Ucn3 neurons in the MePD may be glutamatergic similarly to the Ucn3 neurons in the hypothalamic perifornical area. Glutamate is possibly required to trigger network activity of the MePD Ucn3 neurons similar to the role glutamate plays in the ARC KNDy network (14). This allows the network to reach a certain level of excitability that promotes signalling via Ucn3 to activate CRFR2 expressing GABAergic neurons, which in turn downregulate MePD kiss1 activity, resulting in suppression of GnRH pulse generator activity. Thus, NMDA and AMPA receptor antagonism possibly cancels MePD Ucn3 glutamatergic activation of inhibitory GABAergic CRFR2 expressing inter-neurons preventing the decrease in GnRH pulse generator activity (Fig. 6C). However, fast synaptic activity occurs due to synaptic neurotransmitter release and modulation of this activity is known to be a key target for neuropeptides. Indeed, urocortins can also function endogenously as modulators of excitatory glutamatergic transmission in the amygdala, with *in vitro* studies showing modulation of glutamatergic excitatory post-synaptic potentials via pre- and postsynaptic CRFR2-mediated mechanisms (39). Therefore, we cannot exclude the possibility that Ucn3 activity in the MePD is augmenting glutamatergic transmission to activate downstream inhibitory GABA neurons.

The fact that GABAergic and glutamatergic receptor antagonism within the MePD did not alter LH pulsatility without optogenetic stimulation of Ucn3 neurons is an important factor in our proposed mechanisms and neural circuits within the MePD. This observation indicates that under basal non-stress conditions MePD Ucn3 neurons are relatively quiescent, thus solely pharmacologically blocking the upstream GABA and glutamate inputs to the GABAergic MePD projections without a corresponding increase in Ucn3 activity would make little difference to the net effect of the MePD over the KNDy neuronal system.

Stress elevates Ucn3 mRNA (19) and c-fos expression in the MePD (18) while chemogenetic inhibition of MePD Ucn3 neurons prevents stress-induced suppression of LH pulsatility (3). The MePD sends stress-activated efferents to the PVN (18) and MePD Ucn3 neurons have been shown to project directly to the PVN (21). In the current study, we demonstrate for the first time that optical stimulation of MePD Ucn3 projections in the PVN inhibits pulsatile LH secretion. Recently, chemogenetic activation of PVN CRF neuron perikarya was shown to rapidly suppress LH pulse frequency, however optogenetic activation of their nerve terminals in the ARC does not affect the firing of KNDy neurons (40) and directly injecting CRF in the ARC also has no effect on LH secretion (41), suggesting the involvement of an indirect mechanism inhibiting GnRH pulse generator activity. The MePD has been shown to send Ucn3 neuronal projections to the PVN and receive Ucn3 projections from the PVN, indicating bidirectional communication between these two brain regions via Ucn3 signalling (21). Moreover, the MePD sends direct efferent projections to KNDy neurons of an unknown neurochemical phenotype (8). Therefore, stressful stimuli may be processed by PVN CRF or Ucn3 neurons and this information may be relayed via the MePD Ucn3 neuronal circuitry to the ARC KNDy neurons to modulate their activity (see Fig. 6D). Ultimately, the MePD may function as a novel relay centre between the PVN and ARC to regulate stress signals to the GnRH pulse generator and contribute to the crosstalk between the reproductive and stress axes. Central administration of Ucn3 augments the HPA axis response to restraint stress in rats (19) and increases hypothalamic CRF concentrations and circulating levels of CORT in mice (42). Overexpression of Ucn3 in the hypothalamic perifornical area increases anxiety-like behaviour (38) and activation of the CRFR2 in the lateral septum is involved in stress-induced persistent anxiety in mice (43). In the medial amygdala, Ucn3 neuronal populations are shown to consist of both projection neurons extending to the PVN, the bed nucleus of the stria terminalis and the suprachiasmatic nucleus (21,34) and local Ucn3 interneurons (32,33). We have previously shown chemogenetic inhibition targeting both local and efferent MePD Ucn3 neurons prevents psychosocial stress-induced release of CORT (3). In the present study, we show that selective optogenetic activation of MePD Ucn3 projections in the PVN elevates CORT secretion, thus MePD Ucn3 projections to the PVN are critical for the regulation of CORT secretion. Further work is required to identify whether only the MePD Ucn3 projecting efferents to the PVN are crucial for regulating CORT secretion. The presence of E_2_ is known to up-regulate basal CRF mRNA levels in the PVN in rodents and induce high levels of circulating corticosterone providing a sensitizing effect to the activity of the HPA axis (44,45). Moreover, the CRFR2 promoter contains a classical estrogen receptor response element (46), indicating that CRFR2 activity is upregulated in the presence of E_2_. Therefore, we implanted the OVX mice with E_2_ capsules for HPA axis sensitization in mice similarly to what has been reported in previous studies (44,47–49). However, E_2_ levels may have fallen lower than expected, to low or potentially below the physiological range observed in diestrous mice, as evidenced by little or no effect on LH pulse frequency. However, an effect on LH pulse amplitude and mean LH levels was evident in the E_2_ replaced animals (25). Although there was weak E_2_ negative feedback on LH pulses in our model, our main reason for E_2_ replacement was to sensitizes the HPA axis (44,47–49). PVN-CRF neurons express mRNA transcripts for CRFR2 (50) and CRFR2 primarily activate the Gαs adenylyl cyclase/cAMP signalling pathway (51) indicating that signalling via the CRFR2 located on PVN-CRF neurons may result in activation of this neuronal population and hence the HPA axis (see Fig. 6D). Therefore, Ucn3 released following optical stimulation of MePD Ucn3 projections in the PVN may directly activate the PVN-CRF neurons, signalling via CRFR2, to stimulate CORT release as observed in the present study. However, this hypothesis requires further experimental interrogation.

Selective CRFR2 antagonism has been shown to reverse the stress-induced inhibition of LH pulses (52) where central administration of astressin2-B, a specific CRFR2 antagonist, blocked the restraint-stress induced suppression of pulsatile LH secretion in rats (53). We have shown specific antagonism of CRFR2 in the MePD blocked the inhibitory effects of predator odor on the GnRH pulse generator in mice (3) and activation of MePD Ucn3 neurons delayed puberty in mice (31). Therefore, we know CRFR2 neurons are involved in processing stressful information and coordinating the activity of the GnRH pulse generator accordingly. Additionally, CRF is also implicated in reducing pulsatile LH secretion via CRFR1 expressing neurons in mice (54). Chronic stress levels of CORT decrease LH pulsatility (55) and recently an acute rise in CORT sustained for 30-60 min was shown to inhibit pulsatile LH release, but only in OVX mice with estrogen replacement (56) and after a delay of approximately 30 min following the CORT rise. From our observations, inhibition of LH pulsatility occurs immediately after the stimulation of MePD Ucn3 projections in the PVN in OVX E_2_ replaced mice, thus it is unlikely the result of negative feedback from CORT release but rather a central suppression of GnRH pulse generator activity possibly from the activation of PVN CRF neurons, although the mechanism involved has yet to be established. Moreover, acute psychosocial stress exposure reduces LH pulse frequency in OVX mice (3,57), suggesting E_2_ is not necessary for the inhibition of the GnRH pulse generator in response to psychosocial stress (3,57). Therefore, acute stress pathways possibly act via CORT and E_2_ independent pathways to suppress LH pulses. In our study, optical stimulation of MePD Ucn3 local and efferent neurons is supposed to mimic acute exposure to psychosocial stress as this has been shown to be processed by Ucn3 activity in the MePD, thus supporting the idea of a central suppression of GnRH pulse generator activity mediated by MePD Ucn3 neurons.

The PVN CRF neuronal population was not traditionally considered important for mediating stress-induced suppression of pulsatile LH secretion, as PVN lesion studies have shown that this region is not critical in mediating the stress-induced suppression of the reproductive axis (58). However, with the recent observations showing that chemogenetic activation of PVN CRF neurons indirectly inhibits the GnRH pulse generator it is plausible that the stress neural circuitry includes upstream signalling via the MePD. Therefore, a possible route may involve stressful information processed by the PVN CRF neurons relayed via the MePD Ucn3 neuronal circuitry to the ARC KNDy neurons to modulate their activity (see Fig. 6D). The neighbouring central nucleus of the amygdala is known to act as a relay centre in pain control where the central lateral part receives nociceptive input from the parabrachial nucleus and the central medial part is involved in regulating the output of defensive behaviour and nociception (59). Local and efferent MePD Ucn3 signalling plays a critical dual role in regulating GnRH pulse generator activity and CORT secretion, thus MePD Ucn3 neuronal populations and their projections may represent a novel nodal centre in the interaction between the reproductive and stress axes.

Our findings show for the first time that stimulation of local Ucn3 neurons in the MePD inhibits GnRH pulse generator activity involving an interplay between Ucn3, GABA and glutamate signalling within the MePD. Furthermore, we interrogate the MePD Ucn3 neuron projection terminals in the PVN demonstrating that stimulation of the MePD Ucn3 efferent projections in the PVN elevates CORT secretion to stress levels while suppressing the GnRH pulse generator. How the local neural circuit within the MePD engages with efferent projections from the MePD to the PVN and the interplay between Ucn3, GABA and glutamate neurons to mediate the suppressive effect on LH pulsatility and the stimulatory effect on the HPA axis remains an exciting question for the future.

